# The Set Increment with Limited Views Encoding Ratio (SILVER) Method for Optimizing Radial Sampling of Dynamic MRI

**DOI:** 10.1101/2020.06.25.171017

**Authors:** S. Sophie Schauman, Thomas W. Okell, Mark Chiew

## Abstract

**Purpose:** To present and assess a method for choosing a fixed increment between spokes in radially sampled MRI that results in higher SNR than other radial sampling increments.

**Theory and Methods:** Sampling uniformity contributes to image SNR when reconstructed using linear methods. Thus, for a radial trajectory, uniformly spaced sampling is ideal. However, uniform sampling lacks post-acquisition reconstruction flexibility, which is often needed in dynamic imaging. Golden ratio-based increments are often used for this purpose. The method presented here, Set Increment with Limited Views Encoding Ratio (SILVER), optimizes sampling uniformity when a limited number of temporal resolutions are required. With SILVER, an optimization algorithm finds the angular increment that provides the highest uniformity for a pre-defined set of reconstruction window sizes. SILVER was tested over multiple sets and assessed in terms of uniformity, noise amplification, and SNR both in simulations and in acquisitions of a phantom and healthy volunteers.

**Results:** The proposed algorithm produced trajectories that, for the optimized window sizes, had higher uniformity and lower image noise than golden ratio sampling both in a simulated single-coil system and in a multi-coil system, assessed using simulation, phantom, and in vivo experiments. The noise in SILVER optimized trajectories was comparable to uniformly distributed spokes whilst retaining flexibility in reconstruction at multiple temporal resolutions. In a resting state fMRI experiment, tSNR increases at different spatial and temporal resolutions ranged from 21-72%.

**Conclusion:** SILVER is a simple addition to any sequence currently using golden ratio sampling and significantly increases sampling efficiency and SNR.

## 1. Introduction

Many recent MRI methods, especially highly accelerated and/or dynamic imaging, use radial k-space sampling because of its benefits related to reduced sensitivity to motion (1), and spatially incoherent aliasing when sampled below the Nyquist limit (2).

In some dynamic MR experiments, it is useful to be able to retrospectively reconstruct the same data with different temporal resolutions. Examples of this include generating high temporal resolution navigators for motion correction of lower temporal resolution data (3,4) and retrospective binning of data into frames of unequal length based on respiratory or cardiac phase (5,6). In these cases, the radial golden ratio (GR) method (7) is often used. In the GR method, the direction of each k-space sampling spoke is determined as a set angle increment from the previous spoke, such that each new spoke intersects the largest gap in k-space by the golden ratio. This approach results in relatively high uniformity for any number of subsequently acquired spokes and allows for complete flexibility in reconstruction at multiple temporal resolutions. It also does not repeat the exact same spoke during an experiment, eliminating the risk that retrospectively binned data will contain repeats of the same spoke.

However, the most SNR efficient 2D radial sampling method in terms of minimizing the gaps in k-space for a fixed number of spokes is uniformly distributed spokes. The cost is that uniform radial sampling allows for reconstruction of the data at only a fixed temporal resolution and requires complete a-priori knowledge of how many k-space spokes will be combined to form each frame. It does therefore not have the same flexibility as GR sampling.

In this paper, an alternative method of sampling, which allows for a tradeoff between the flexibility of GR and the SNR efficiency of uniformly distributed spokes, is presented. The idea behind this method is that GR is often unnecessarily general for imaging experiments that will be reconstructed at only a small number of temporal resolutions. The proposed method uses numerical optimization of the angular increment between subsequent spokes for a restricted set of window sizes (number of spokes per frame) with the aim of maximizing the uniformity of sampling within that set. By relaxing the requirement for ‘near-uniformity’ to only apply to a specific set of window sizes, it was hypothesized that higher sampling efficiency could be achieved whilst maintaining the favorable properties provided by the GR methods. In this paper a procedure for choosing the fixed angular increment between subsequent spokes is presented and the resulting sampling patterns compared with both the GR method and with radially uniform sampling in simulations, phantoms, and in vivo. We call the proposed method the Set Increment with Limited Views Encoding Ratio (SILVER) method.

## 2. Theory

### 2.1. Properties of set increment sampling

In this study radial sampling trajectories with a pre-set, or fixed, increment between spokes are considered. Set increment sampling is easy to design and implement as one only needs to decide on a single angle increment, *θ*, that is added to the projection angle of the previous spoke. For radial spokes that span the full diameter of a circular k-space, *θ* can be defined using a sampling ratio, 0 < *α* < 1, such that *θ* = *α* × 180°.

When α is a rational fraction, the exact same spoke will eventually be repeated, whereas if the step is irrational, or rational with a very large denominator in its simplest form (practically irrational), no two spokes acquired within the duration of the experiment will be the same. If multiple temporal resolutions are required, it is beneficial to use irrational sampling to avoid duplicate spokes in the different window sizes, especially if data is combined across all frames to construct a temporal average image, for example for coil sensitivity map estimation. Also, in imaging experiments where the length of a frame is unknown a-priori, practically irrational increments are preferred to avoid acquiring the same spoke again within each frame. Setting *α* to the reciprocal golden ratio 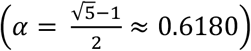 which is often referred to as the most irrational number (8), is therefore commonly done in order to sample without repeats.

Regardless of whether α is rational or irrational, acquiring data in k-space with a set angular increment generates data that can be flexibly reconstructed, as any grouping of *N* subsequently acquired spokes will have the same k-space sampling pattern, only rotated (Figure 1). This allows for sliding window (9) and view sharing (10) reconstructions.

**Figure 1.**
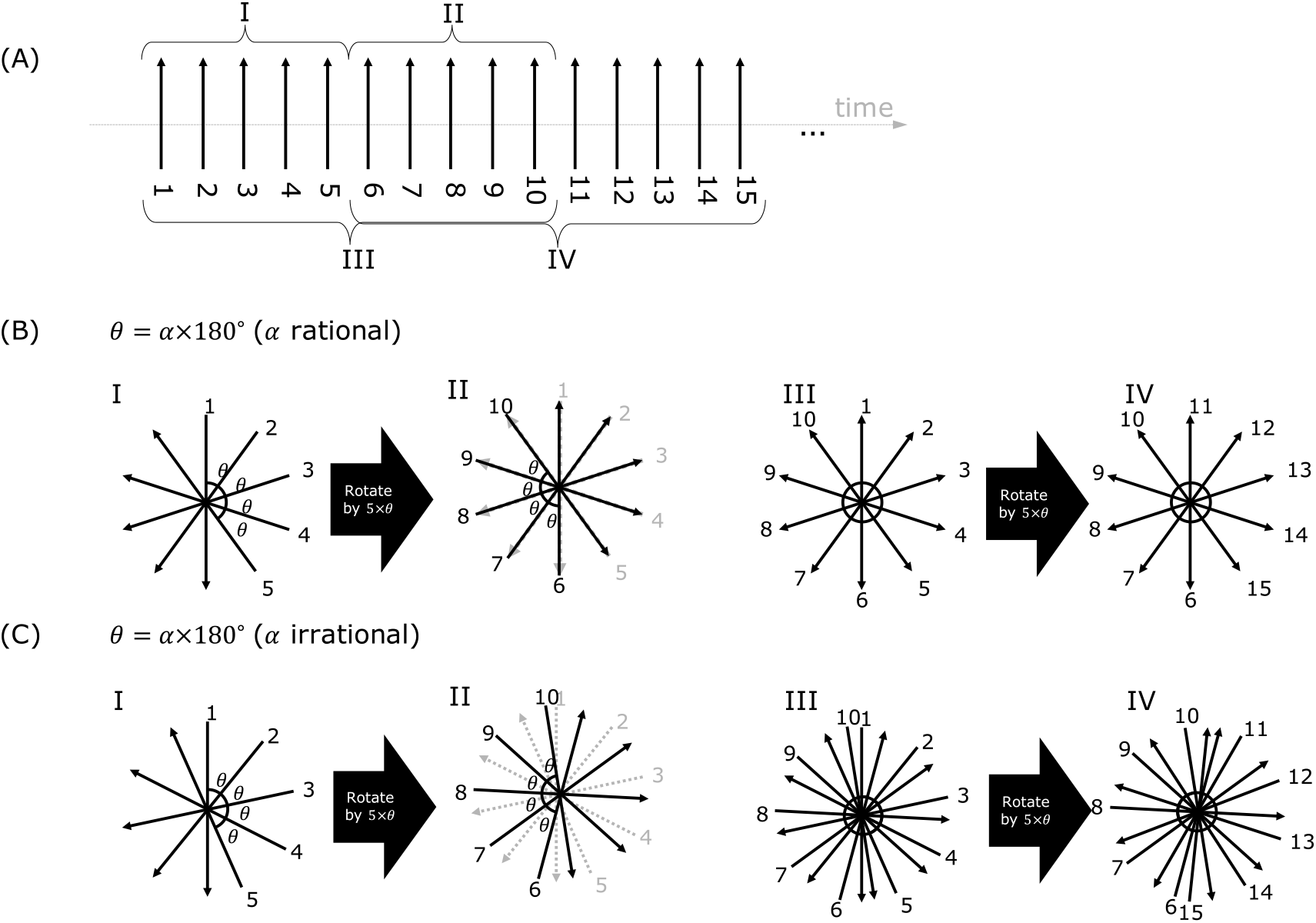
The effect of set increment radial sampling with rational and irrational increments. (A) demonstrates four different groupings of subsequent spokes (I, II, III, and IV) and gives each spoke a number. (B) shows how the subsequent spokes in the different frames relate if α is rational (in this case 1/5), showing how both non-overlapping and overlapping frames have repeats of the same spokes. (C) shows the effect of an irrational α; Non-overlapping subsequent frames have no repeat spokes, and overlapping frames have some of the same spokes. In both (B) and (C) a subsequent frame (whether overlapping or not) is simply a rotaion of the previous frame and will thus have the same level of uniformity in each case.

In this study, all sampling patterns that are studied use set increment sampling. Uniform radial sampling with full width spokes can be achieved with set increment sampling by choosing the angular increment between subsequently acquired spokes to be 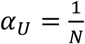, where *N* is the number of spokes used to reconstruct one frame. In GR sampling, on the other hand, the step is instead approximately 111.24°, with 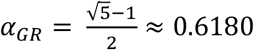. The aim of SILVER is thus to find a more optimal increment, 0 < α < 1, when near uniformity is only required for a certain set of window sizes.

### 2.2. Sampling and SNR

With non-uniform sampling density, denser areas of sampling (e.g. near the centre of k-space or where two radial sampling spokes are nearly overlapping) have to be down-weighted. If each sample is correctly weighted the signal can be accurately reconstructed. However, noise measured in each sample is averaged with noise from all other samples when using a Fourier transform, and noise averaging is most effective with equal weighting of each noise sample. Sampling with non-uniform density thus leads to non-ideal averaging, and noise amplification in MRI (11,12). The proposed method thus aims to improve uniformity of sampling to improve SNR.

Another way of considering the relationship between sampling and SNR is to consider the linear model governing the acquisition and reconstruction methods. When sampling k-space in a non-uniform manner, off the Cartesian grid, reconstruction generally involves taking the pseudo-inverse of the acquisition operator, ***E***, either directly or through iterative methods. ***E*** is defined by the forward model of MRI, ***y*** = ***Ex*** + ***n***, where ***y*** is the data recorded by the scanner, ***x*** is the object that is being imaged and ***n*** is thermal noise. The operator ***E*** is generally composed of a coil sensitivity operator that models how each receive channel weights the object ***x*** spatially, and a sampled Fourier transform that models how the data is acquired using spatial encoding along a specific trajectory. In a pseudo-inverse reconstruction (***E***’***E***)^-1^ must be calculated or iteratively estimated, and its diagonal entries (assuming pre-whitened noise covariance) will govern noise amplification (13). Therefore, if the spokes are not well distributed, some of the columns/rows of ***E*** will be similar, leading to poor conditioning of ***E*** and greater noise amplification. Strictly speaking, the noise amplification depends on the interactions between both the sampling trajectory and the coil sensitivities. However, other sampling methods, such as CAIPI (14), have been shown to reduce noise amplification robustly without requiring consideration of specific coil sensitivity profiles.

### 2.3. Measuring sampling efficiency

Because SNR intrinsically depends on the uniformity of the local k-space sampling density, different methods for estimating the local sampling density have been proposed. For 2D radial sampling in particular, Winkelmann et al. defined sampling density by the inverse of the average azimuthal distance between adjacent spokes (7). For spiral imaging the sampling density has been estimated using the speed of spiraling around k-space (15), or using voronoi cells (16). For 3D radial sampling numerically defined approaches using voronoi cells on spheres have also been used (17). An alternative to these methods of considering density distributions, specific to radial sampling, is to consider a physical model based on electrostatic force or potential between charges placed on the spoke tips. This constitutes a version of the Thomson problem (how charges would naturally be distributed on a sphere with minimal potential energy) where each charge has a pair on the opposite side of the sphere (13,14). Methods like these are extensively used to choose diffusion sampling directions in diffusion tensor imaging <u>(20,21)</u>. Electrostatic methods, although more commonly used in 3D sampling, can also be easily applied to 2D sampling problems. An electrostatic potential minimizing model was used in this version of SILVER because of its easy extension to 3D in the future, and its high penalty for overlapping spokes.

In the electrostatic potential model, the radial sampling pattern is treated as an ensemble of unit charges placed on both ends of each spoke spanning the diameter of the unit circle. The total energy of the system is thus defined as:

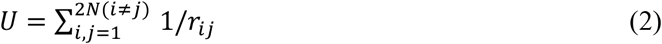

Where *r*_*ij*_ is the distance between the i^th^ and j^th^ points, and *N* is the number of spokes (2*N* is therefore the number of spoke tips).

Sampling efficiency compared to radially uniform sampling, *η*, was then defined as the ratio of total electrostatic potential stored in the system of point charges, *U*, to a system with the same number of spokes in the lowest possible energy state (uniformly distributed spokes), *U*_*ref*_:

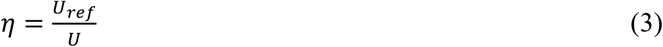

*U*_*ref*_ is included in the efficiency metric, *η*, as a normalizing factor, so that comparison of efficiencies at different *N* is possible. For sampling with a set increment, *U* reduces to a function of only α, and the number of spokes, *N*. For a uniform radial distribution α = 1/*N*, so *U*_*ref*_ is simply a function of *N*.

## 3. Methods

### 3.1. Optimization

The SILVER method was formulated as an optimization problem: to maximize the minimum efficiency, *η*, for a pre-defined set of window sizes, *S = {N*_*1*_, *N*_*2*_, *N*_*3*_, *…}*. The objective function was therefore defined as:

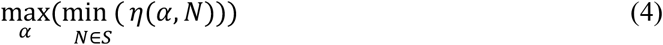

Where *N* is the number of spokes in the window, *α* is the fixed sampling ratio (as defined in section 2.1), and *η* is the electrostatic potential based efficiency metric (defined in section 2.3). The optimization was performed in MATLAB R2018b (The MathWorks, Inc., Natick, Massachusetts, United States) using optimcon in the Optimization Toolbox using the interior-point minimization algorithm. To avoid local minima the optimization algorithm was restarted 100 times with 99 initial values of *α* drawn from a uniform probability distribution between 0 and 1 and one run with the golden ratio as the starting value. A flowchart of the main SILVER optimization method is shown in Figure 2a.

**Figure 2.**
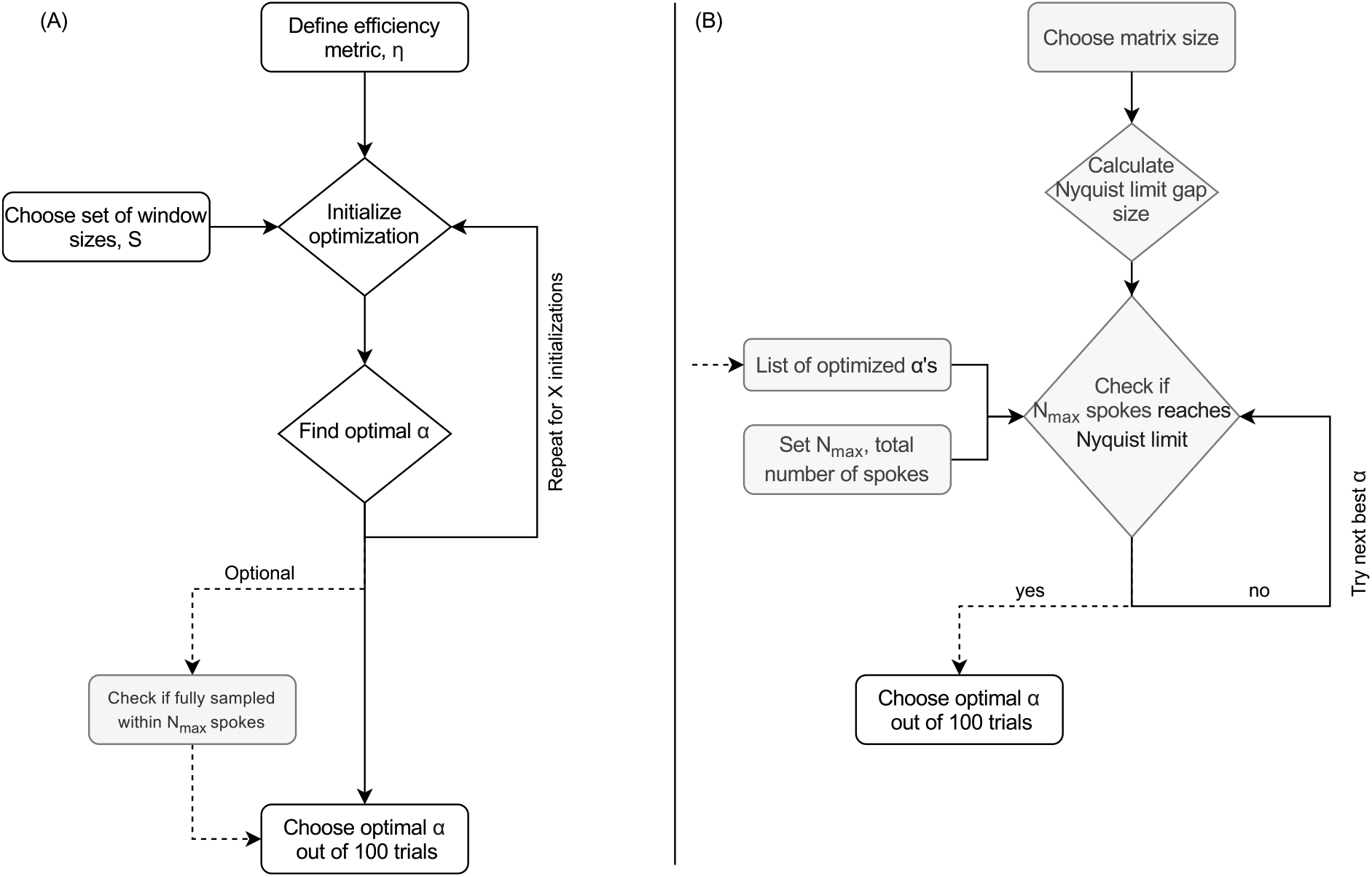
(A) shows the main flowchart of the SILVER algorithm. Because the cost function is non-convex, multiple repeated initializations are used. In our experiments, the number of initializations, X, was 100. (B) shows the optional extra final step which ensures that the trajectory does not start to repeat spokes within the full length of the experiment and eventually reaches the Nyquist limit, such that by combining all spokes in an experiment one can estimate the coil sensitivity maps from the data itself.

An additional, optional, criterion was included in the optimization algorithm to check for Nyquist coverage at some maximum number of spokes to enable coil sensitivity estimation from a fully sampled, time averaged, set of k-space data. A flowchart for this additional step in the algorithm is shown in Figure 2b.

First, the effect of optimizing for multiple discrete temporal resolutions was examined. The efficiency for sets with *S =* {*M*, 2*M*}, and S = {M, 2*M*, 3*M*} were studied, with *M* set to 4, 16, and 32. As a special case, *S* was chosen to consist of five Fibonacci numbers (*S* = {5, 8, 13, 21, 34}), the window sizes where GR sampling performs optimally.

Continuous ranges of window sizes were also explored to determine over what range of window sizes SILVER can perform better than GR. This matters for scans where the data is binned retrospectively (e.g., for cardiac or respiratory gating), and the number of consecutive spokes allocated to each bin is thus unknown at the time of acquisition, but a small continuous range of possible window sizes can be pre-determined. These sets contained a minimum window size of *M* spokes, and all intermediate window sizes up to a maximum window size, *M+R*, such that *S* = {*M, M+1*, …, *M+R*}. *M* was set to 8, 16, and 32, and *R* was set to 3, 5 and 7.

### 3.2. Simulating and measuring noise amplification

To compare SILVER optimized imaging more directly with GR and uniform radial sampling, noise amplification in images acquired with the three methods was predicted using Monte Carlo simulations and compared to the predicted efficiency using the uniformity metric, *η*. Gaussian complex noise was generated and reconstructed linearly using an iterative conjugate gradient reconstruction for the trajectory being examined. The reconstruction algorithm was implemented in MATLAB using the built-in preconditioned conjugate gradients method, pcg, with 100 iterations. A final filtering step was performed which involved setting the unsampled corners of (Cartesian) k-space to zero to avoid signal in regions unconstrained by the circular sampling window. This was done by taking the final image output from the iterative reconstruction, applying a Cartesian fast Fourier transform, using a circular mask to set the corners of k-space to zero and then transforming back to image space using a 2D inverse fast Fourier transform. The reconstruction was run 100 times for the same trajectory with different instances of noise, and the resulting pixelwise standard deviation across instances was measured. The resulting noise standard deviation maps were compared using a Wilcoxon signed-rank test at a significance level of 0.005 to control for multiple comparisons.

First, the trajectories were trialed in a simplified system containing a single coil with uniform sensitivity. The linear forward operator, ***E***, was thus defined by the trajectory alone. An oversampled acquisition with 60 spokes for a matrix size of 30 x 30, starting at 0° from the x-axis, and then incremented by *θ* = α × 180° for each subsequent spoke, was simulated. A SILVER angle was optimized for three sets: the sets, *S*_1_ = {50, 60}, *S*_2_ = {60, 70}, and *S*_3_ = {58, 60, 62}. The increment for radially uniform sampling was 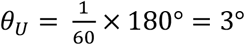, for GR sampling: *θ*_*GR*_ = 111.246°, and for the three versions of SILVER: *θ*_*SILVER,1*_= 147.561°, *θ*_*SILVER,2*_= 48.840°, *θ*_*SILVER,3*_ = 71.412°.

Next, the effect of trajectory choice with a fixed set of coils was explored to determine whether optimizing for the trajectory alone can reduce noise amplification in a more complex parallel imaging setting. The same simulation as for the single coil case was performed, but with 10 spokes and eight coil sensitivity maps (compressed from a 32-channel phantom measurement using singular value decomposition (22)). The uniform sampling angle was thus 18°, and SILVER was optimized for *S*_1_ = {5, 10}, *S*_2_ = {10, 15}, and *S*_3_ = {9, 10, 11}, giving optimized angles 32.759°, 126.936°, and 98.680°, respectively. Additionally, the experiment was repeated 179 more times with the starting angle shifted by 1°. This was to further study how the trajectory and coil sensitivity maps interacted. The noise means of each angle for each trajectory were also compared using the Wilcoxon signed-rank test at a significance level of 0.005 to control for multiple comparisons.

### 3.3. Example use case – TURBINE fMRI

After the simulation experiments, SILVER was assessed in both in a phantom and in vivo in two healthy volunteers. The in vivo data was acquired with approval by the local ethics committee.

SILVER was applied to a multi-shot 3D fMRI experiment, with a 3D radial-Cartesian sequence consisting of rotating EPI blades, TURBINE (4). In multi-shot 3D fMRI experiments, there are SNR gains to be made relative to 2D multi-slice methods, but the acquisitions are more sensitive to motion. Therefore, reconstructing a fast navigator can be useful for estimating motion which can be used to correct the full reconstruction. SILVER can thus be used to optimize TURBINE to two temporal resolutions simultaneously: the fast navigator and the full imaging temporal resolution.

Each TURBINE-EPI blade was fully sampled along the phase encoding direction (z-direction), which allowed 2D radial k-space data to be generated for each slice separately, after Nyquist ghost correction and an inverse Fourier transform along the z-direction. The sequence parameters were TE = 28 ms, TR = 50 ms, FA = 15°, FoV read (rotating in the x-y plane) = 200 mm, and FoV phase (along the z-axis) = 16 mm (phantom)/32 mm (in-vivo). The base resolution was 2 mm isotropic. Two experiments, with two temporal resolutions each, were performed. The two fast navigator temporal resolutions were 400 ms (8 blades per frame), and 500 ms (10 blades per frame). The two imaging temporal resolutions were 2.3 s (46 blades per frame) and 2.75 s (55 blades per frame). Each slice (slice thickness 2 mm) was reconstructed on a matrix size of 30 x 30 with an in-plane spatial resolution of 6.7 x 6.7 mm^2^ for the navigator, and a matrix size of 100 x 100 with an in-plane spatial resolution of 2 x 2 mm^2^ for the fMRI imaging. These temporal resolutions were chosen to assess SILVER at both Fibonacci numbers (8 and 55 blades per frame), which is optimal for GR, and non-Fibonacci numbers (10 and 46 blades per frame). For comparison, uniformly sampled data was acquired in four acquisitions with α_U_ = 1/8, /10, 1/46, and 1/55 respectively. GR sampled data was acquired in a single acquisition with α_GR_ = 0.6180. SILVER was optimized for two sets S_Fib_ = {8, 55}, and S_no-Fib_ = {10,46} with α_SILVER,Fib_ = 0.87279 and α_SILVER,no-Fib_ = 0.29786. The scan time for each trial was 1 min 22 s and each one was repeated two times to assist in noise characterization. These scans were acquired to assess noise performance across a wide range of parameters both in-vivo and in a phantom, with repeats used to subtract the signal and focus only on the noise to compare with simulations. The same significance tests as for simulation data were performed on the phantom and in vivo data using the different trajectories but at a significance level 0.0042 to account for the 12 tests performed in each case.

In vivo, two additional, more realistic, resting state fMRI scans were acquired without repeats, with α_SILVER,no-Fib_, and α_GR_, for 5 min 6 s each. For these, tSNR was directly compared between GR and SILVER. Again, statistical significance was measured using a Wilcoxon signed-rank test at a significance level 0.025.

The data were reconstructed slice-by-slice after Nyquist ghost correction, global phase correction, and an inverse Fourier transform across the fully sampled phase-encoding direction (superior-inferior), with the same iterative reconstruction method as had been used for the Monte-Carlo noise simulations described in section 3.2. The short acquisitions that had two repeated acquisitions were phase aligned by multiplying the k-space data from the first acquisition by a global phase factor, *e*^*iφ*^, on a spoke-by-spoke basis, which minimizes the L2 distance between the spokes (complex data vectors) in the first and second acquisition. A complex subtraction of the repeats then yielded pure noise images that were studied by taking the pixel-wise standard deviations to measure noise levels. Additional predictions of noise were performed for the measured coil sensitivities and coil covariances. In this case, noise was not only predicted for the first *N* spokes, but simulated noise data was reconstructed using continuously sampled spokes leading to a rotating trajectory that varies frame-by-frame, such that the expected temporal noise could be estimated as the standard deviation across time.

The longer resting state fMRI scans were reconstructed using the same linear reconstruction framework, but only a single repeat was acquired. This was done to demonstrate the tSNR benefit even in the presence of signal fluctuations. These data were examined by measuring the pixelwise tSNR taking the temporal mean divided by the temporal standard deviation. Additional compressed sensing reconstructions were performed, with sparsity enforced in the spatial wavelet domain (4-level Daubechies db2) to assess whether the benefit from SILVER persisted for non-linear reconstructions. The compressed sensing reconstruction replaced the iterative conjugate gradient method with the fast iterative shrinkage thresholding algorithm (FISTA (23)) with a step size of 10^−4^ for low resolution reconstruction with 10 spokes/frame and 10^−6^ for high resolution reconstruction with 46 spokes/frame. The regularisation factor was set to 10^−6^, and the number of iterations was 50. These parameters were chosen empirically.

The coil sensitivity maps for all reconstructions were estimated using the adaptive combine method, using coil images reconstructed from all the data from the GR acquisition, and used for all reconstructions. The SILVER acquisitions also produced fully sampled temporal mean images when combined across all frames that could have been used for coil sensitivity estimation. However, the GR sensitivity map was used for all to reduce sources of variance in the results.

## 4. Results

The SILVER optimization succeeded in improving the minimum efficiency within all the targeted sets compared to GR. As shown in Figure 3a, for the trialed sets, the improvement varied from 0.067% (for SILVER optimized for five Fibonacci numbers) to 4.2% (for SILVER optimized for the set S = {4, 8}). 13 out of the 16 trialed sets achieved improvements larger than 1%. Figure 3b shows an example of efficiencies for different window sizes sampled with GR and two of the SILVER optimized increments. Within the targeted window sizes, the two SILVER optimizations consistently outperformed GR, even reaching higher efficiency than GR at its peaks at Fibonacci numbers.

**Figure 3.**
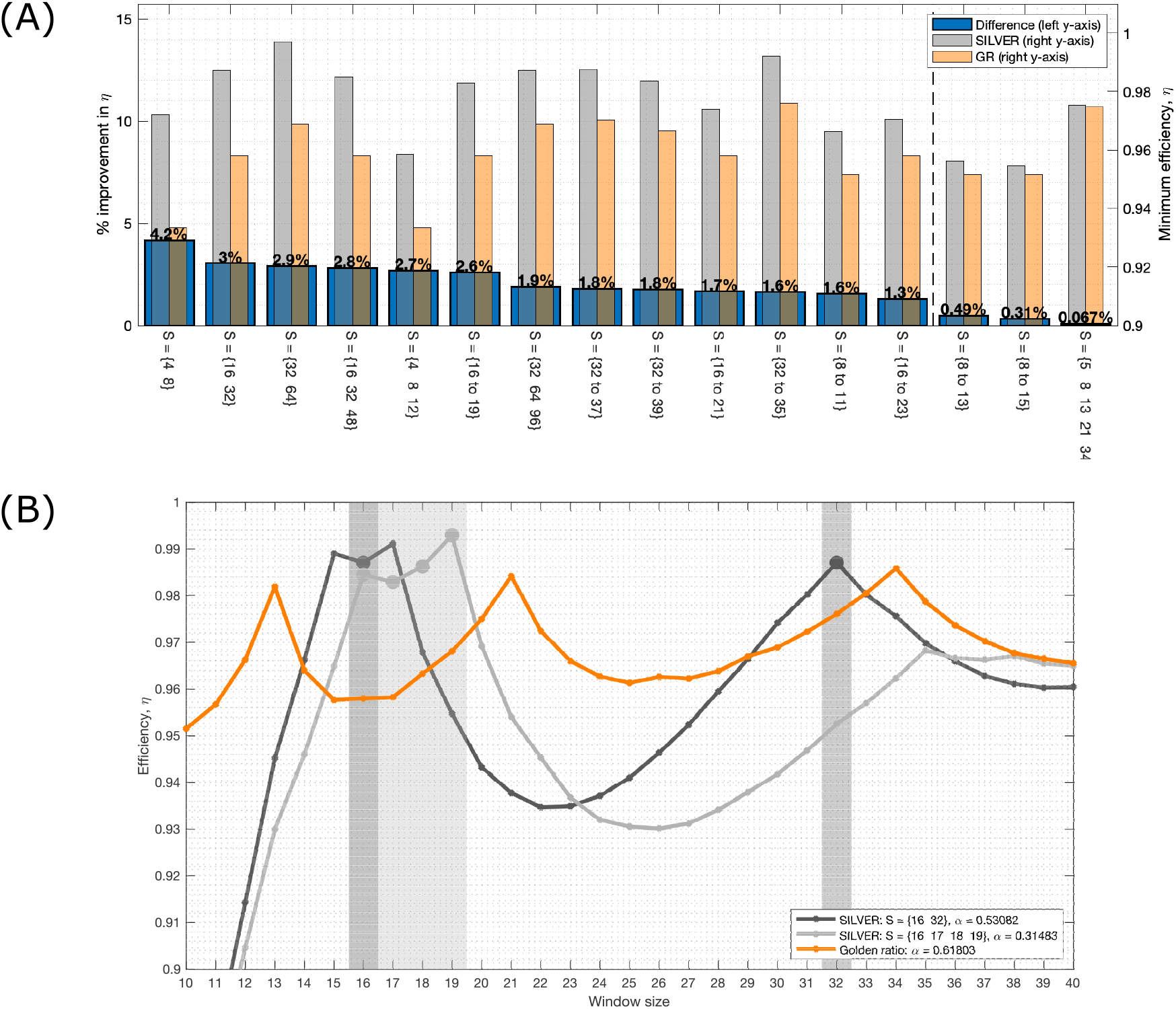
(A) shows the result of the SILVER optimization for 16 sets of window sizes ordered by percentage improvement in sampling efficiency for the worst window size within the set, measured using the electrostatic potential metric, *η*. The vertical dashed line separates the sets where the improvement was more or less than 1%. (B) shows the efficiency of two of the SILVER optimizations (*S* = {16, 32} and *S* = {16 to 19}) along with GR in more detail. The shaded areas with round markers show the window sizes that the two SILVER increments were optimized for respectively. Both optimizations reach higher efficiency for their optimized window sizes than GR, which has characteristic peaks at Fibonacci numbers 13, 21, and 34.

Noise levels in the single coil simulations (Figure 4) followed the predicted order with lowest noise in uniformly sampled data and highest noise in GR sampled data, with the SILVER optimized trajectories having noise levels in between GR and uniform. All differences were statistically significant except the two SILVER trajectories optimized for S = {50, 60}, and S = {60, 70}. In single coil simulations with other angle increments noise levels correlated with efficiency estimated using the electrostatic potential metric, especially near optimal efficiency (Supporting Information Figure S1).

**Figure 4.**
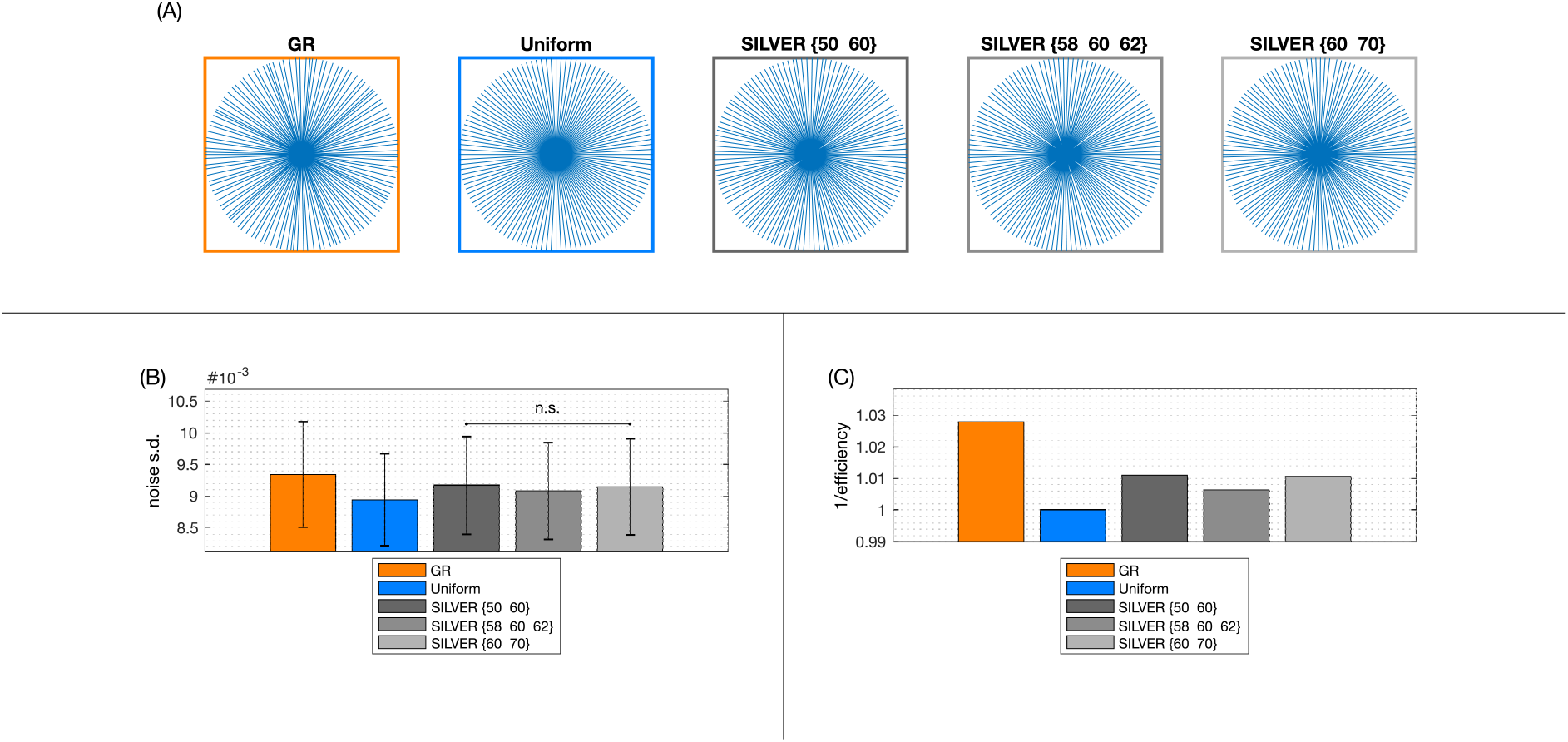
(A) shows the five 60-spoke trajectories that were compared in the single coil simulations. (B) shows the average noise standard deviation measured for each trajectory. The error bar shows the standard deviation across space. (C) shows the inverse of the efficiency measured using the electrostatic potential metric. It is correlated with the result in (B) as expected.

In multi-coil simulations (Figure 5), the same trend was observed, with highest noise in GR and lowest in the uniform sampling case. However, as the trajectories interacts with the coils, the relative order of SILVER sets did not directly correspond to the estimates based on efficiency of the trajectory alone. The effect of the trajectory interacting with the coil sensitivities was also noticeable when the trajectory was rotated with respect to the coils (Figure 6), although these fluctuations were of smaller magnitude than the differences between the methods. One of the SILVER sets, S = {5, 10}, produced noise that on average was similar to GR with a non-significant difference, whereas the other two sets had significantly lower average noise.

**Figure 5.**
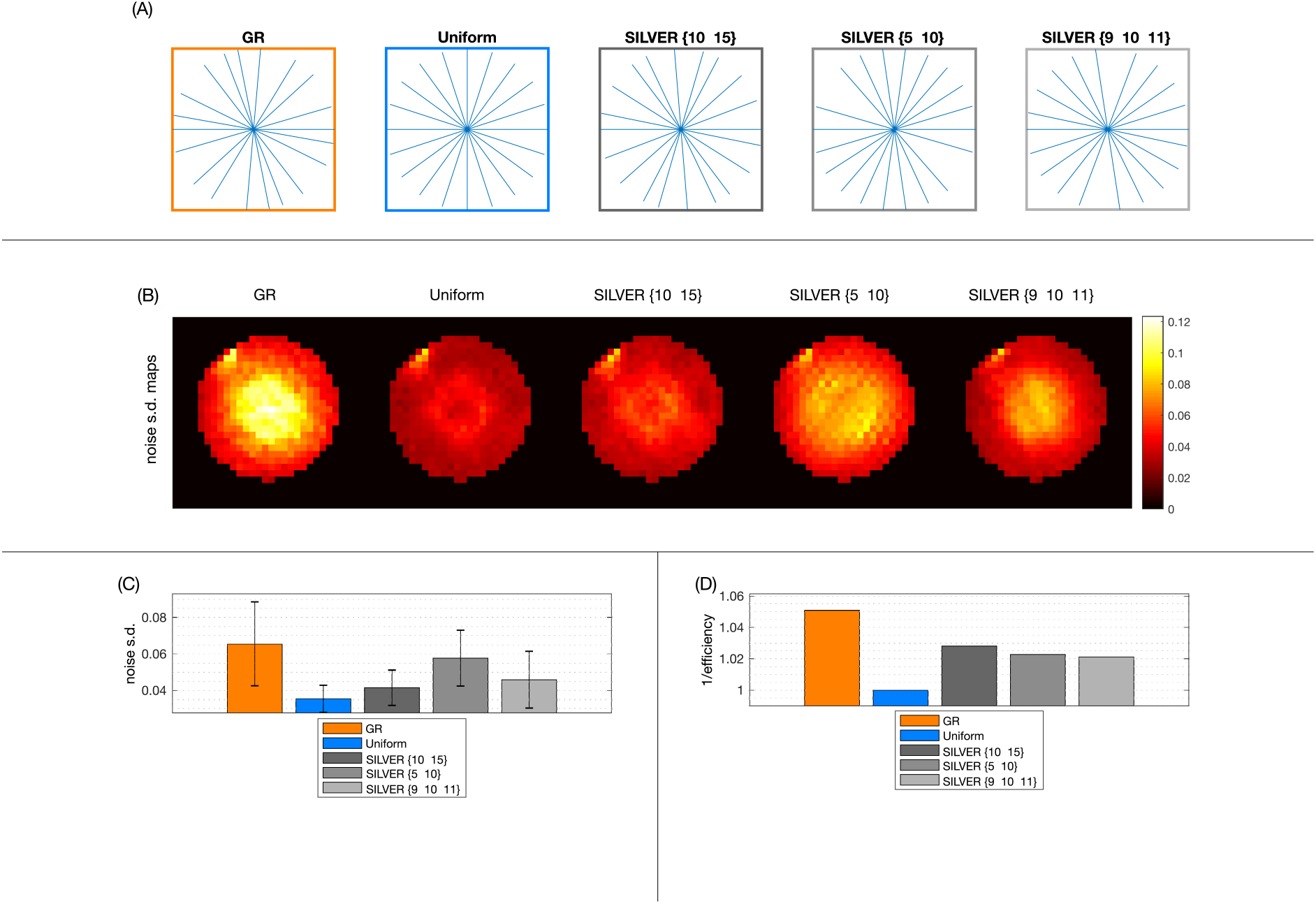
(A) shows the five 10-spoke trajectories used in multi-coil simulations. (B) shows the resulting noise maps for each trajectory. The noise level is calculated by taking the standard deviation across noise initializations. (C) shows the average noise standard deviation measured for each trajectory. The error bar shows the standard deviation across space. All SILVER trajectories produced noise that was significantly lower than GR but higher than uniform sampling. (D) shows the inverse of the efficiency measured using the electrostatic potential metric. It is not as correlated with the noise s.d. (C) as it was for single coil data (Figure 4

**Figure 6.**
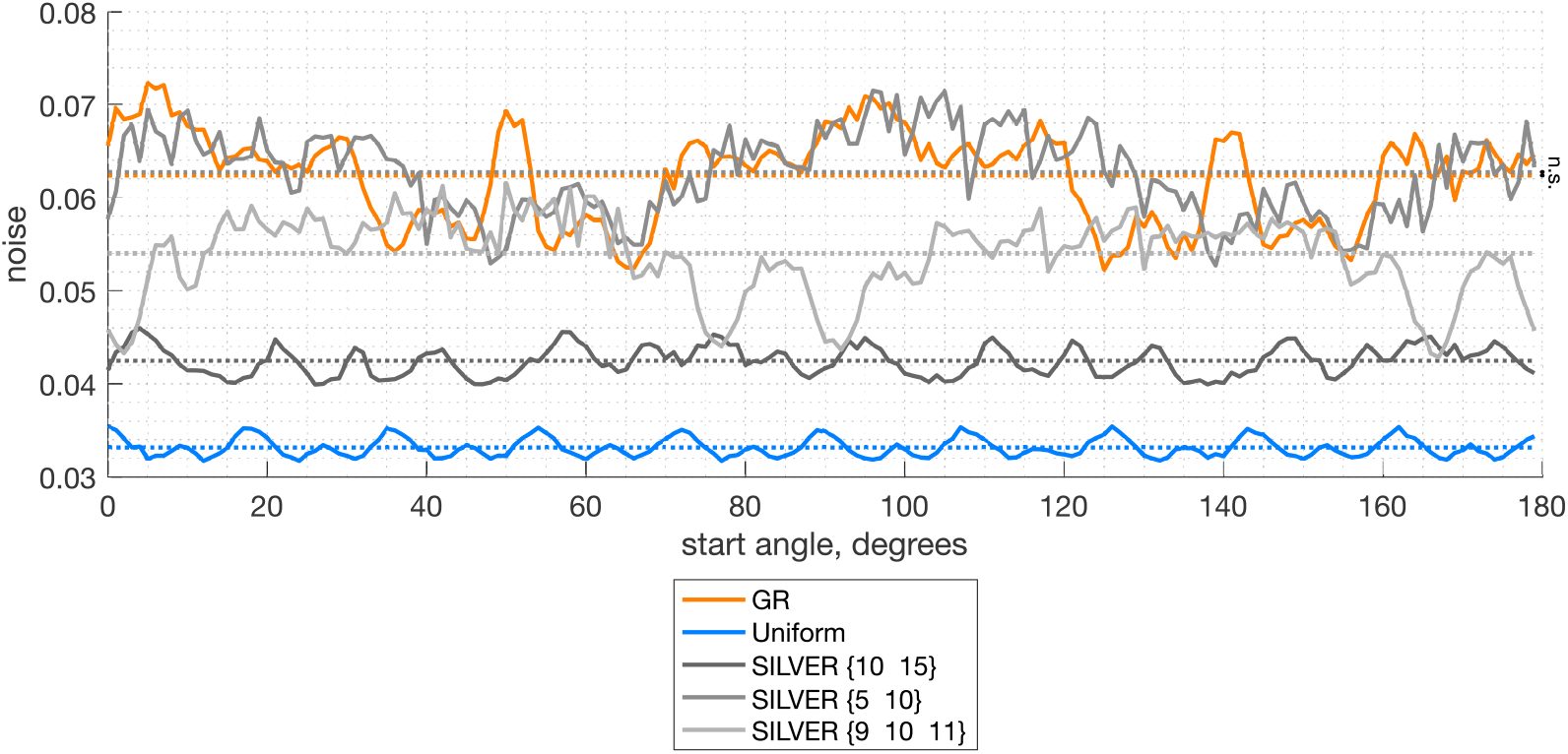
Average noise level for the trajectories shown in Figure 5a as the trajectory is rotated with respect to the coil sensitivity maps. Uniform sampling has an expected period of 18 degrees because that is the spoke spacing.

The acquired phantom data corresponded closely to the noise predicted from the coil sensitivities, noise covariance, and the performed trajectory, shown in Figure 7. For all four window sizes, SILVER had a performance with noise levels lower than GR. The difference in median noise level between uniform and GR sampling was +26%, +48%, +25%, and +5%, for window sizes 8, 10, 46, and 55. The difference between uniform and SILVER was much smaller: 0%, -1%, +9%, and +1% for the same window sizes. The noise maps, normalized to the uniform sampling median value, are shown in Figure 7a, and the corresponding violin plots of measured and predicted noise values shown in 7b.

**Figure 1.**
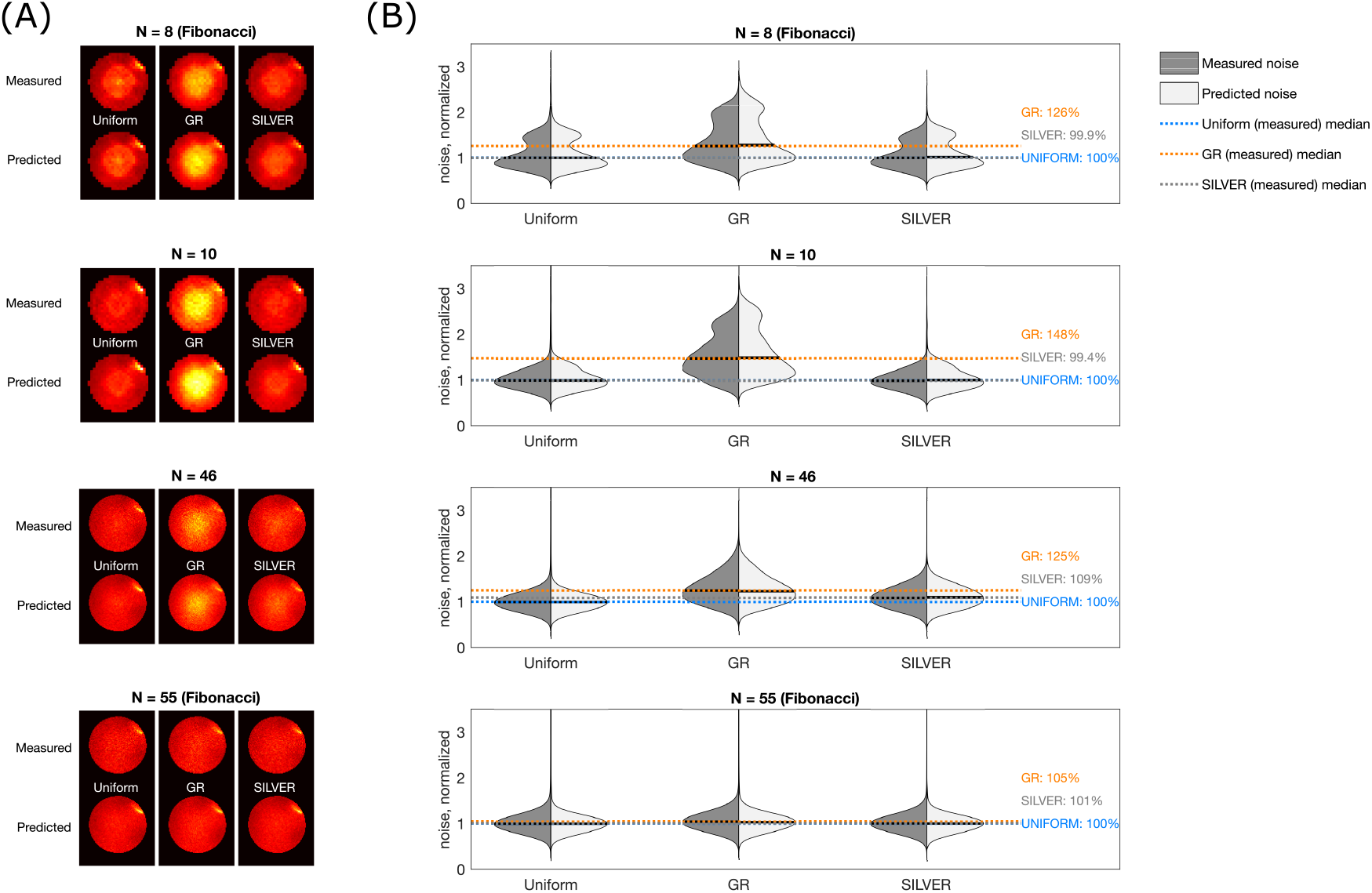
Noise maps from simulations and phantom measurements match closely: (A) shows the noise maps (normalized to the uniform noise median) and (B) shows the same data in violin plots. The medians of the measured data are shown in dashed (blue = uniform, gray = SILVER, orange = GR).

Similar results to the phantom results were found in vivo (Figure 8). However, a small discrepancy between predicted and measured noise was found due to additional signal fluctuations arising from physiological noise and any residual aliasing. The difference in median noise level between uniform and GR sampling for subject B shown in Figure 8 was in this case +27%, +48%, +18%, and -2%, for window sizes 8, 10, 46, and 55. The difference between uniform and SILVER was +4%, +22%, +11%, and -1% for the same window sizes. Similar results were measured in the other subject (Supporting Information Figure S2).

**Figure 8.**
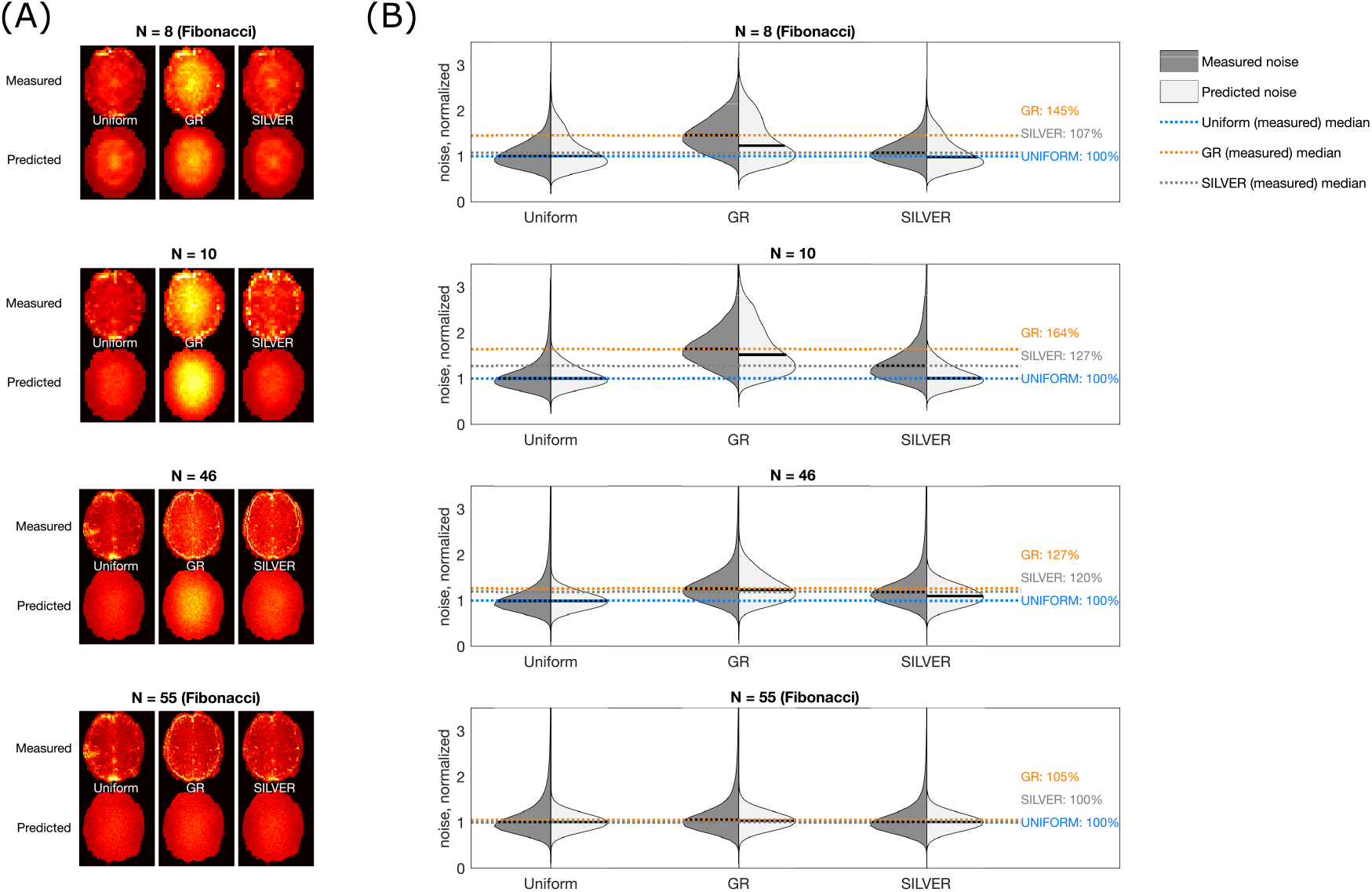
Noise maps from simulations and in vivo measurements match closely, (A) shows the noise maps (normalized to the uniform noise median) and (B) shows the same data in violin plots. The medians of the measured data are shown in dashed (blue = uniform, gray = SILVER, orange = GR). The localised noise increases around the sagittal sinus in the measured data are probably due to pulsatility.

The longer fMRI experiment also showed improvements in tSNR for SILVER compared to GR. Using a linear reconstruction, the tSNR for the fast navigator (10 spokes/frame) improved from 10.46 to 17.98 (72%) by using SILVER instead of GR, and for the imaging resolution (46 spokes/frame) the tSNR improved from 16.40 to 19.83 (21%). Figure 9 shows tSNR maps in an example slice from subject A and tSNR histograms across all slices. The results in subject B were comparable (Supporting Information Figure S3).

**Figure 9.**
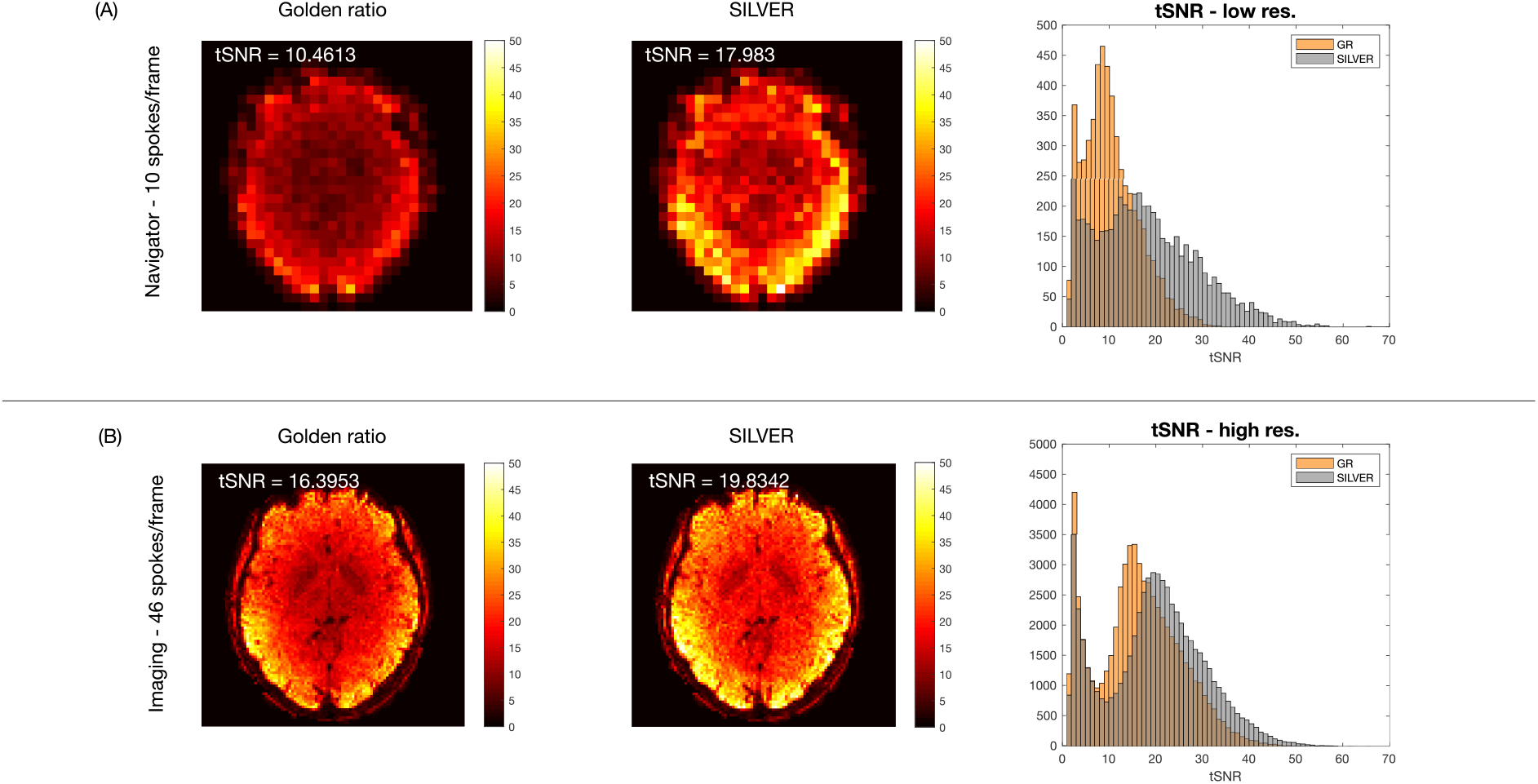
tSNR rises significantly by using SILVER instead of GR both when the data is reconstructed with 10 spokes/frame (A), and when reconstructed with 46 spokes/frame (B) using linear parallel imaging. An example slice is shown in the images with the average tSNR calculated across all slices shown on top. The histograms show the data from all slices combined.

Also, when using a more sophisticated reconstruction (wavelet regularized compressed sensing), SILVER continued to demonstrate higher tSNR than GR (Figure 10 shows an example slice of subject A). The difference was somewhat smaller than in the linear case (35% improvement for the low-resolution navigator and 15% for high-resolution imaging), but nonetheless substantial. The results from subject B were similar and are attached in the Supporting Information (Figure S4).

**Figure 10.**
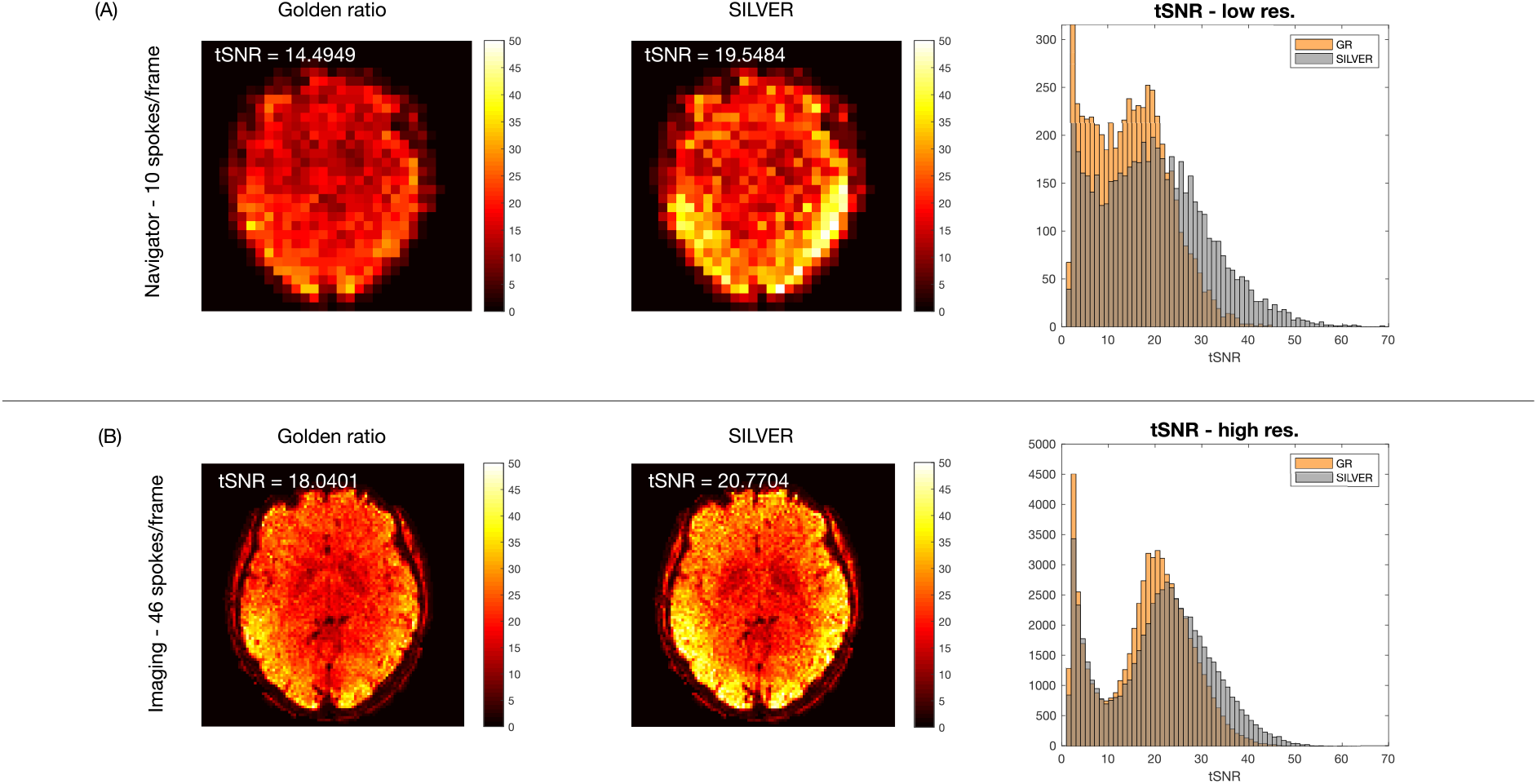
tSNR rises significantly by using SILVER instead of GR both when the data is reconstructed with 10 spokes/frame (A), and when reconstructed with 46 spokes/frame (B) using non-linear compressed sensing reconstruction. An example slice is shown in the images with the average tSNR calculated across all slices shown on top. The histograms show the data from all slices combined.

Images of the temporal average and temporal standard deviations for both linear and non-linear reconstructions in both subjects are attached in Supporting Information Figures S5-S8.

## 5. Discussion and conclusions

We have presented a method of choosing an optimal angular increment for dynamic radial MRI when the set of window sizes to consider is constrained. We have shown that this method results in decreased noise compared to GR sampling with only a minor change required to the acquisition protocol: the single value that controls the angular increment.

We showed that small differences in uniformity measured using the electrostatic potential metric can lead to large differences in the amount of thermal noise in radial acquisitions. The results showed a max difference of ∼4% in electrostatic potential, but up to ∼70% difference in tSNR. The largest improvements in uniformity were for small sets including only two temporal resolutions, and in these cases improvements over GR at Fibonacci numbers was even possible. For larger sets, the improvement in uniformity was lower. However, even when no significant improvement was achieved, SILVER converged onto a trajectory with the same efficiency as GR (i.e. GR or a tiny golden angle (24)). So, there was no disadvantage to optimizing for SILVER-based increments on any set of window sizes of interest. SILVER is easy and fast to implement (for example, SILVER for S = {10, 46} took ∼7 s to run on a MacBook Pro, 2.7 GHz Intel Core i7, without any optimization for speed) and will only be needed to be optimized once for every set of window sizes of interest.

SILVER is optimized without knowledge of the coil sensitivities, and the trajectory inevitably interacts with the sensitivity maps in a parallel imaging reconstruction. By trialling SILVER at different starting angles, the effect of this interaction could be assessed. With uniform sampling, the same trajectory with respect to the coils are acquired for each frame, but with GR or SILVER each frame has a rotated version of the trajectory and can thus have varying noise amplification in different frames. The temporal behaviour and periodicity for these methods should be studied further for applications where the temporal signal characteristics can affect image analysis, such as in fMRI.

The results showed good consistency between predicted and measured noise, especially in the phantom, but slight deviations were observed in vivo, likely due to signal changes originating in pulsatility and subtle motion. In the data presented, no motion correction was applied since the subjects were well behaved and not using motion correction avoids skewing tSNR measurements due to effects originating in the interpolation. SILVER showed an improvement in tSNR for both linear and non-linear reconstructions indicating that the method is not limited by a specific reconstruction approach.

The example application of SILVER presented in this paper was a multi-temporal resolution fMRI experiment, where one resolution was intended to be used as a navigator, the other as the imaging resolution. Examples of other applications of SILVER include retrospective gating for cardiac/respiratory motion, including XD-GRASP (25), or multiple temporal resolutions for simultaneous angiography and perfusion imaging (e.g. CAPRIA(26)).

Because conditioning of the acquisition operator depends on the coil sensitivities, as well as the sampling trajectory, optimizing for sampling uniformity alone cannot be guaranteed to always produce the optimal trajectory leading to the lowest noise. However, without knowledge of the specific coil sensitivity maps to be used, optimizing for uniformity alone is a tractable step in the right direction, similar to how in CAIPI (14) the conditioning of the parallel imaging problem is improved by changing the trajectory alone. In this paper, we have shown that optimizing the trajectory without knowledge of the coils in general leads to reduced noise as coils intended for parallel imaging are independently optimized by the coil manufacturers. In one case SILVER was observed to produce on average slightly higher noise than GR (when varying starting angles were trialed, Figure 6), but the difference was non-significant. Similarly, in the noise measurements for 55 spokes in-vivo and 8 spokes in the phantom (Figures 7 and 8), GR and/or SILVER produced lower noise than uniform sampling, which could be due to interaction with the coil sensitivities or simply due to variance in the measurements. Again, in these cases, the size of these effects was very small (less than 2% difference in noise standard deviation).

In future work, the SILVER framework could also be extended to optimize for properties other than trajectory uniformity. This could be done through design of alternative cost functions, including cost functions that consider the temporal incoherence of the operator for temporally regularized reconstructions. Additional constraints on the optimization could also be included to limit the SILVER angles to be small for less eddy current contamination, an issue that is currently addressed in GR using the tiny golden angle approach proposed by Wundrak et al. (24). However, unlike other work that optimize trajectories more generally (e.g. SPARKLING (27)), SILVER is focused on achieving as large improvements to image quality as possible with as small a change as possible to existing sequences for ease of adoption.

Both SILVER and GR only optimize for contiguously binned spokes, and there is no guarantee that binned data from multiple repeats of radial acquisitions are distributed optimally. Suggestions for overcoming this problem have been proposed for golden ratio sampling when the number of repeats are known a-priori by e.g. Fyrdahl et al. (28) and Song et al. (29), and these methods are equally applicable to SILVER.

Radial sampling does not have to be performed with a fixed increment and could instead be acquired with some other scheme for determining the spoke directions. However, this project focuses on fixed increment radial imaging because of the easy implementation, the consistency in the uniformity of the trajectory across frames, and the ability to use a sliding window for reconstruction if desired. MR sequences that currently use uniform or golden ratio increments could easily be modified to SILVER sampling by changing a single parameter, the increment angle, in the sequence implementation. The SILVER framework also has a potential to be extended to imaging with other trajectories that have previously been derived from the golden ratio, e.g. spirals (30,31), cones (32), and even Cartesian sampling (33).

In 3D, a set increment is harder to define than in 2D. Chan et al. (17) proposed the multi-dimensional golden means method, a 3D analogy of the golden ratio method with a constant azimuthal angle increment and a constant z-axis increment of the tip of the spoke constrained to the surface of a sphere. However, subsequent frames generated with this method are not simply a rotation of the spokes from the previous frame, and therefore not all image frames are guaranteed to have the same sampling efficiency. Because of the increased complexity of 3D radial sampling and the ambiguity of how to define a set increment, we constrained the scope of this paper to focus on the more commonly used 2D case, although extension of SILVER to 3D trajectories, where the electrostatic potential metric of uniformity should work as well, will be considered in future work.

In conclusion, SILVER is a method that generalizes set increment sampling beyond golden ratio derived methods when some knowledge of the number of spokes to combine in a frame is known a-priori, as is commonly the case. Therefore, sometimes SILVER is better than gold, and when it is not, it is just as good.

## 6. Data availability statement

Code for SILVER optimization, all reconstructions, and analysis used in this paper are available on https://github.com/SophieSchau/SILVER. Anonymized data are available on http://doi.org/10.5281/zenodo.4743420 (phantom), http://doi.org/10.5281/zenodo.4743418 (subject A), and http://doi.org/10.5281/zenodo.4743764 (subject B).

## 7. Acknowledgements

This study was supported by funding from the Engineering and Physical Sciences Research Council (EPSRC) and Medical Research Council (MRC) (EP/L016052/1), the Royal Academy of Engineering (RF201617\16\23, RF/132). The Wellcome Centre for Integrative Neuroimaging is supported by core funding from the Wellcome Trust (203139/Z/16/Z). T.O. is supported by a Sir Henry Dale Fellowship jointly funded by the Wellcome Trust and the Royal Society (220204/Z/20/Z).

## Notes

https://github.com/SophieSchau/SILVER

http://doi.org/10.5281/zenodo.4743420

http://doi.org/10.5281/zenodo.4743418

http://doi.org/10.5281/zenodo.4743764

### Competing Interest Statement

All authors have a pending US patent application on the SILVER technique described in this paper.

### Summary of Updates

The paper has been completely overhauled after being rejected with invitation to resubmit during peer-review. The paper has been extended and now includes a new application of the SILVER approach (fMRI instead of ASL) and more detailed phantom and in vivo studies as opposed to the focus on simulation that the first version had.

