## Supporting Information for "The Set Increment with Limited Views Encoding Ratio (SILVER) Method for Optimizing Radial Sampling of Dynamic MRI"

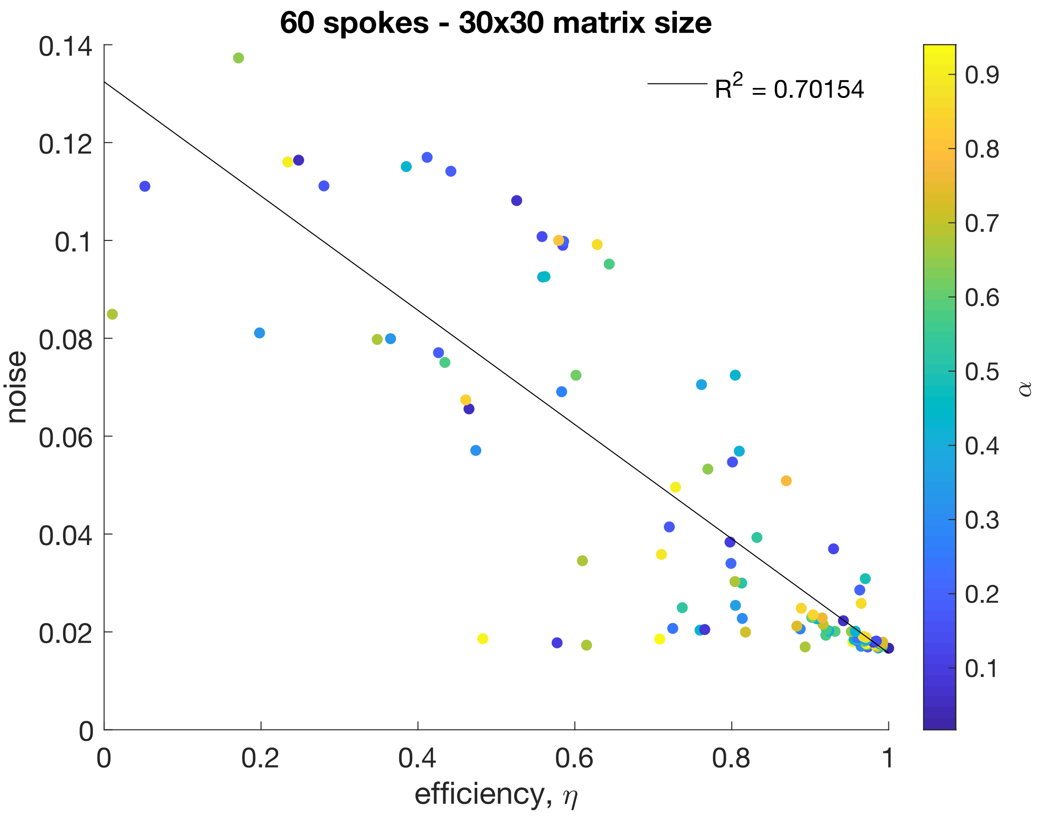


Figure S1 – Efficiencies of 60 spoke trajectories with different set increments, α, correlates with the noise in the reconstructed images. The correlation is stronger when efficiency is closer to 1 probably due to more reliable reconstruction when no large gaps are left in k-space.

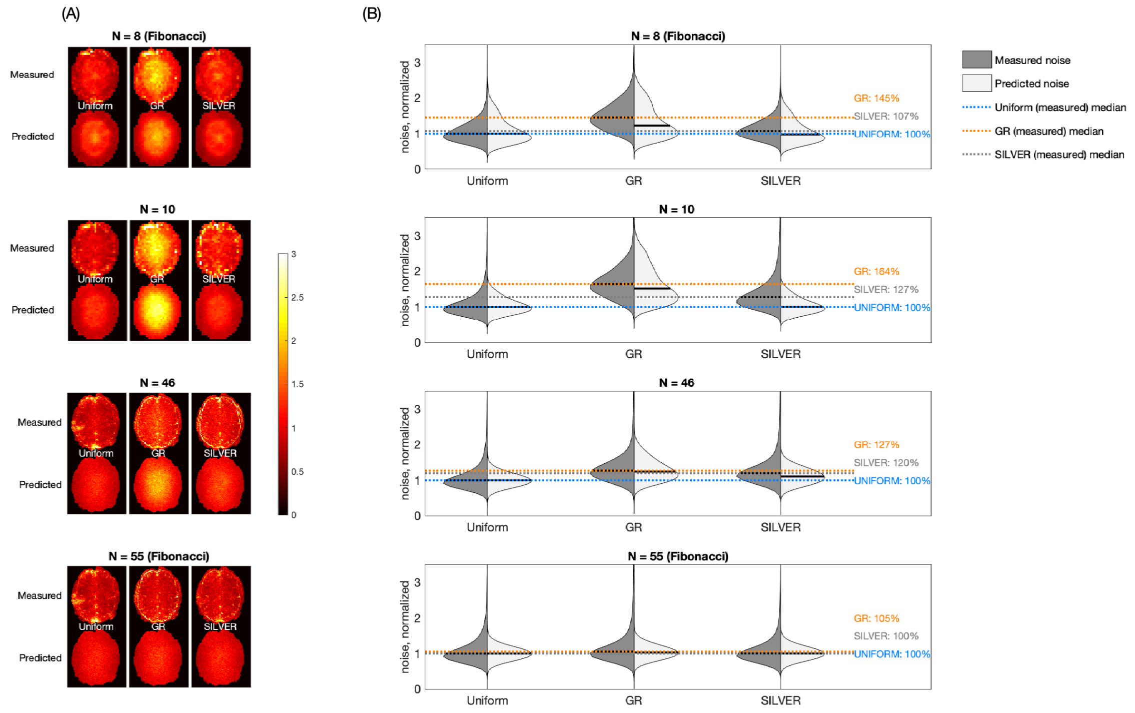


Figure S2 – In vivo noise measurement results for subject A. Equivalent to Figure 8.

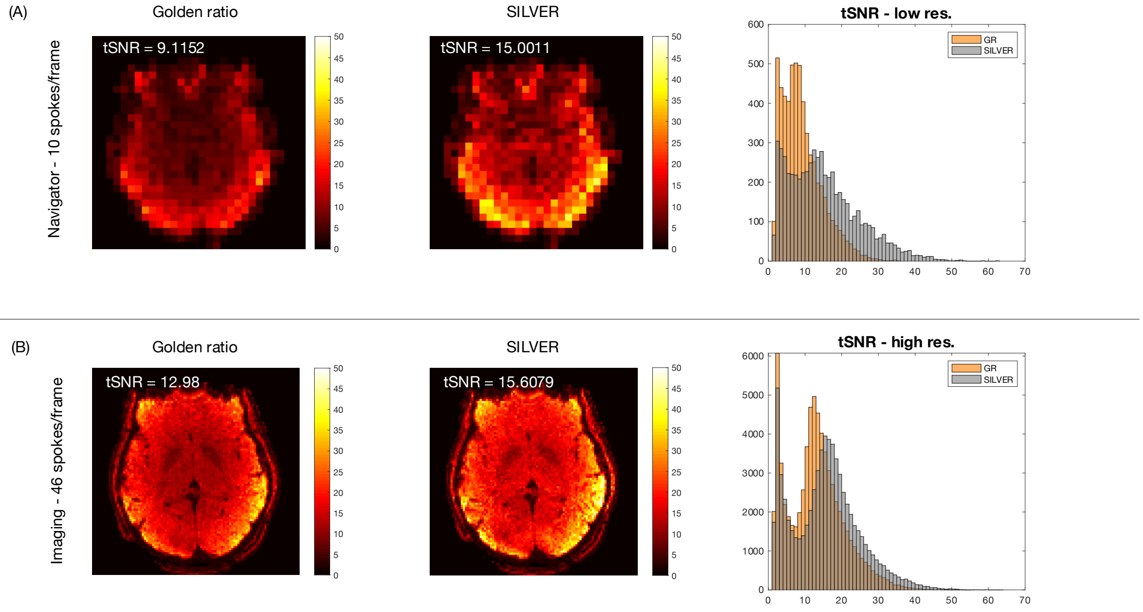


Figure S3 – In vivo tSNR measurements for linearly reconstructed subject B. Equivalent to Figure 9.

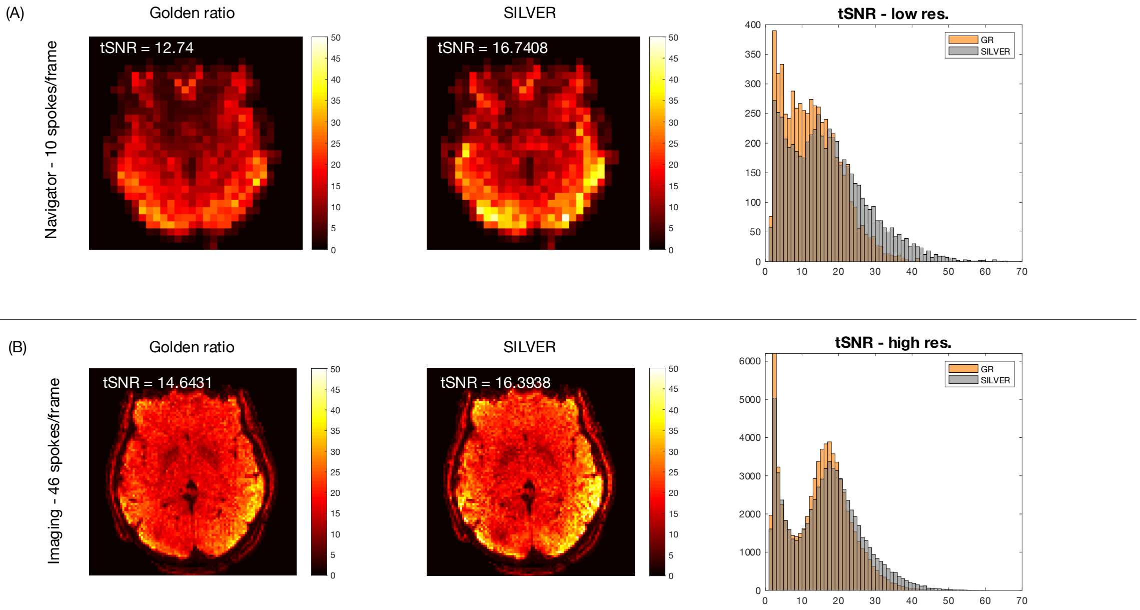


Figure S4 – In vivo tSNR measurements for non-linearly reconstructed subject B. Equivalent to Figure 10.

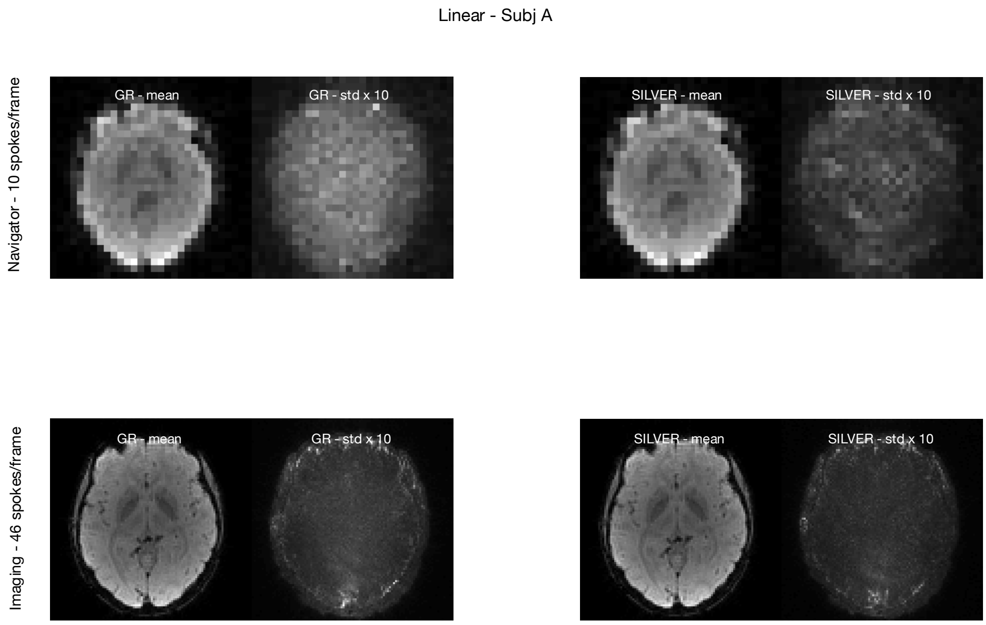


Figure S5– Linearly reconstructed images – Subject A.

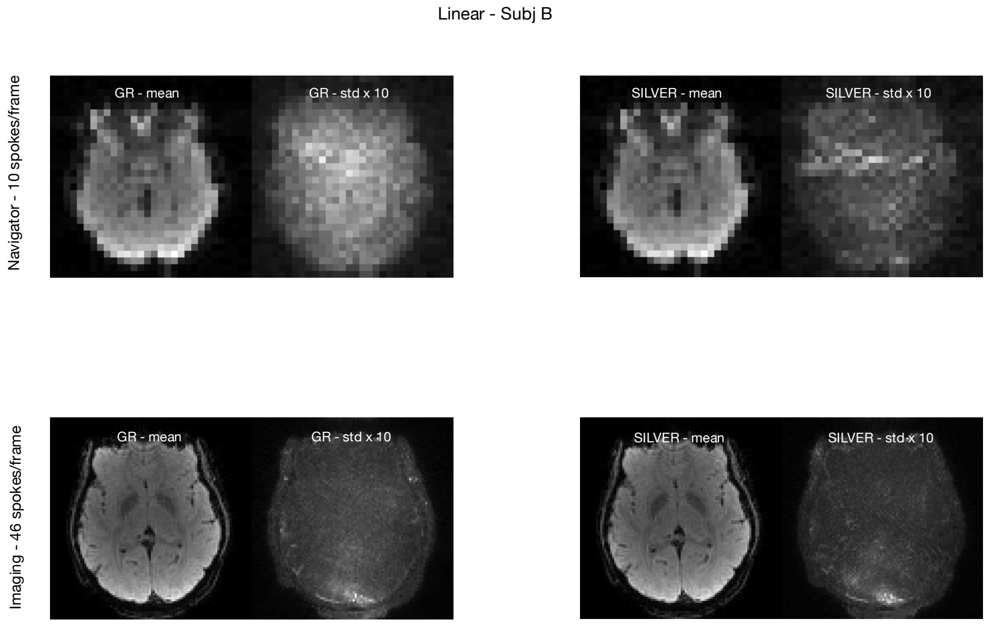


Figure S6 – Linearly reconstructed images – Subject B.

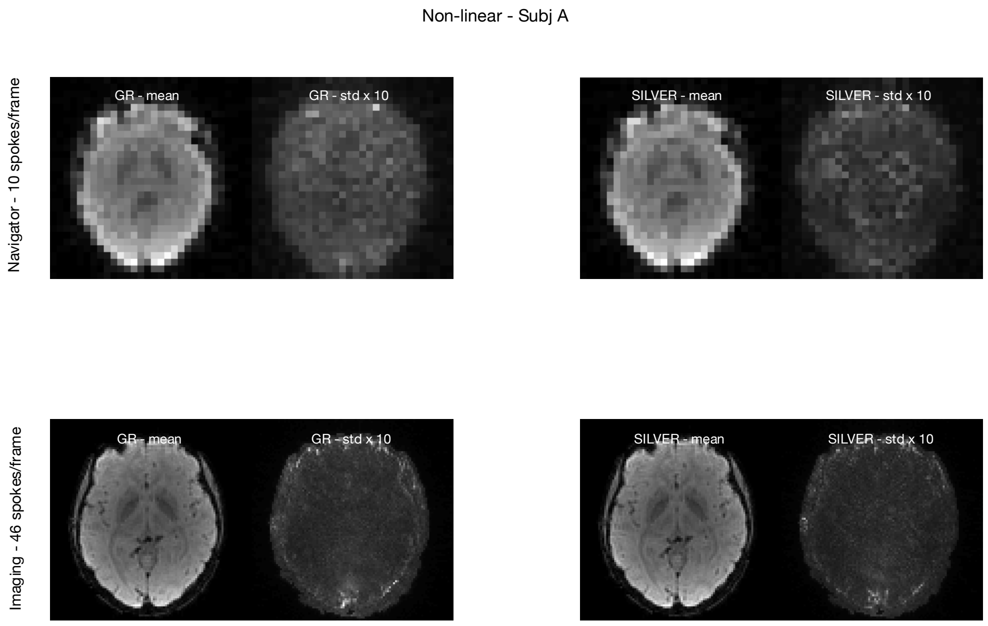


Figure S7 – Non-linearly reconstructed images – Subject A.

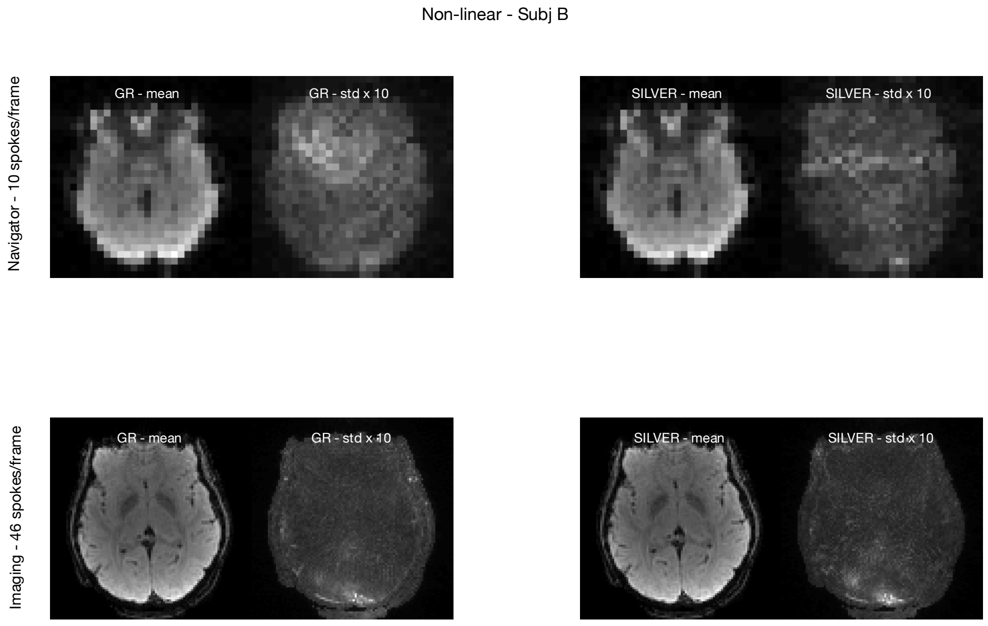


Figure S8 – Non-linearly reconstructed images – Subject B.
